# Two-Dimensional-Dwell-Time Analysis of Ion Channel Gating using High Performance Computing Clusters

**DOI:** 10.1101/2022.09.01.506168

**Authors:** Efthymios Oikonomou, Thomas Gruber, Achanta Ravi Chandra, Sarina Höller, Christian Alzheimer, Gerhard Wellein, Tobias Huth

## Abstract

The power of single-channel patch-clamp recordings is widely acknowledged among ion channel enthusiasts. The method allows observing the action of a single protein complex in real time and hence the deduction of the underlying conformational changes in the ion channel protein. Commonly, recordings are modeled using hidden Markov chains, connecting open and closed states in the experimental data with protein conformations. The rates between states denote transition probabilities that, for instance, could be modified by membrane voltage or ligand binding. Preferably, the time resolution of recordings should be in the range of microseconds or below, potentially bridging Molecular Dynamic simulations and experimental patch-clamp data. Modeling algorithms have to deal with limited recording bandwidth and a very noisy background. It was previously shown that the fit of 2-Dimensional-Dwell-Time histograms (2D-fit) with simulations is very robust in that regard. Errors introduced by the low-pass filter or noise cancel out to a certain degree when comparing experimental and simulated data. In addition, the topology of models, that is, the chain of open and closed states could be inferred from 2D-histograms. However, the 2D-fit was never applied to its full potential. The reason was the extremely time-consuming and unreliable fitting process, due to the stochastic variability in the simulations. We have now solved both issues by introducing a Message Passing Interface (MPI) allowing massive parallel computing on a High Performance Computing (HPC) cluster and obtaining ensemble solutions. With the ensembles, we have optimized the fit algorithm and demonstrated how important the ranked solutions are for difficult tasks related to a noisy background, fast gating events beyond the corner frequency of the low-pass filter and topology estimation of the underlying Markov model. The fit can reliably extract events down to a signal-to-noise ratio of one and rates up to ten times higher than the filter frequency. It is even possible to identify equivalent Markov topologies. Finally, we have shown that, by combining the objective function of the 2D-fit with the deviation of the current amplitude distributions automatic determination of the current level of the conducting state is possible. It is even possible to infer the level with an apparent current reduction due to the application of the low-pass filter. Making use of an HPC cluster, the power of 2D-Dwell-Time analysis can be used to its fullest, allowing extraction of the matching Markov model from a time series with minor input of the experimenter. Additionally, we add the benefit of estimating the reliability of the results by generating ensemble solutions.

## Introduction

Single-channel patch-clamp recordings provide the opportunity to observe a single protein complex during its work with high temporal resolution. That is, observing its transitions from closed to open states and vice versa, termed gating. Single-channel time series contain the most detailed description of gating behavior, which can be obtained with electrophysiological methods. Scientists strive to extract the data hidden in single-channel recordings since the beginning of patch-clamp recordings. The technique to resolve single-channel activity was developed and refined mainly in the 1970s and 1980s (Neher and Sakmann, 1976; Hamill et al., 1981; Benndorf, 1995). A common method to model single-channel kinetics is using Markov Models (Colquhoun and Hawkes, 1977). They are a probabilistic tool that allows approximation of single-molecule behavior. The Markov formalism is essentially an extension of the classical chemical reaction mechanism, where a state corresponds to a stable conformation of the molecule and the rate constants determine the probability of transitions to different conformations. A major challenge is the limited observability of the system. The information about hidden states can be extracted from statistical distributions of the durations of observable events (for an in-depth review see Qin, 2007). An often used and relatively simple way to extract the desired data from a time series with only one active channel is to idealize the time series and to calculate one-dimensional (1D) dwell-time histograms. By applying an exponential fit to the dwell-time histogram, time constants are estimated and then used to infer an underlying model relating kinetics and protein function. Alternatively, a Markov model could be directly estimated from the idealized time series using maximum likelihood estimates (Horn and Lange, 1983; Qin et al., 1997; Colquhoun et al., 2003). 1D-dwell-time histograms do not allow for an unambiguous inference on the underlying Markov scheme, since the connection of states is not preserved. Therefore, the analysis of twod-imensional dwell-time histograms (2D-histograms) was an important step to advance model discrimination (Magleby and Weiss, 1990a). Combining 2D-histograms and simulations of time series further improved analysis (Magleby and Weiss, 1990b; Huth et al., 2006, 2008). With the application of 2D-histograms, connecting adjacent closed and open states, no information is lost and, in theory, this approach should enable the full resolution of the underlying Markov model. Hidden Markov modelling (HMM) approaches are considered the most advanced algorithms for interpreting single-channel gating (Fredkin and Rice, 1992; Albertsen and Hansen, 1994; Qin et al., 2000; Venkataramanan and Sigworth, 2002). They do not require idealization and estimate the likelihood for a given Markov model directly from the time series. The main specific disadvantages of these algorithms are the relative sensitivity to noise (Huth et al., 2006) and temporal limitations imposed by the required low-pass filtering in patch-clamp recordings (Huth et al., 2006; Qin, 2007). The latter one could be a severe issue, since in recent years, the detection of very fast gating kinetics gained increasing attention. The availability of single-molecule data from optical techniques and from molecular dynamics simulations on the timescale of μs makes it possible to correlate data with single-channel patch-clamp recordings. One method to analyze fast current fluctuations is the investigation of β-distributions. If the bandwidth of the recording system is not sufficient to resolve single gating events, the current distribution becomes distinct from a sum of Gaussian distributions and turns into a sum of β-distributions. The deviation from a Gaussian distribution can be used to deduce time constants beyond the filter frequency (for a comprehensive review see Schroeder, 2015). For instance, utilizing β-distributions, very fast gating of viral K^+^ channels (Rauh et al., 2017) and of ryanodine receptors (Hartel et al., 2018) could be resolved. Again, fitting of β-distributions is limited by noise and not sensitive to slow kinetics at all, which introduces problems with inferring complete Markov models from recorded time series. The latter problem was improved by combining an HMM estimator with the fit of β-distributions (Schröder et al., 2005). A general issue with all algorithms mentioned above is to determine how well the fitted model approximates a given time series.

Generally, we consider the current state of single-channel recordings and analysis stagnant because of the tremendous effort involved in thorough modelling. On the other hand, macroscopic recordings commonly used for model generation do not contain the complete kinetic properties of ion channels and the outcome is likely ambiguous. In this work, we addressed the fundamental issues associated with previous versions of the Two-Dimensional Dwell-Time-Fit (2D-fit) (Magleby and Weiss, 1990b; Huth et al., 2006). Therefore, we can now provide a universal framework to extract all the information from single-channel recordings. First off, we improved the computationally very time-consuming task of simulating all the required time series for convergence by implementing parallel computing as previously proposed (Magleby and Weiss, 1990b). Importantly, this approach now allows conducting ensembles of fits for a single problem. Ensembles could then be statistically analysed and used to assess the quality of the results and compare different fit scenarios. With this powerful tool at hand, we optimized the settings of the algorithm. Additionally, we show in this work that ensemble fits are the key to distinguish global fit results and are mandatory for topology discrimination of Markov models. We have implemented and tested two new features. As for the direct HMM estimator (Schröder et al., 2005) the 2D-fit can now be combined with an amplitude histogram fit. Determining the current level in case of apparent reduction due to fast gating (Huth et al., 2006) can now be conducted simultaneously with rate constant estimation. In summary, with massive parallel computing at hand the reliable, and as we consider complete, analysis of a set of times series could be conducted within days or even hours.

## Methods

### Two-Dimensional Dwell-Time fit - overview

The two-dimensional dwell-time fit (2D-fit) with simulations was first introduced by Magleby and co-workers (Magleby and Weiss, 1990b). It was implemented in the KielPatch Program and further refined (Huth et al., 2006). A flow chart of the 2D-fit is illustrated in Fig. 1A. A two-dimensional dwell-time histogram (2D-histogram) is created from a given time series (experimental time series *x_exp_*[*t*], e.g., Fig. 1B) by first idealizing with a jump detector (Fig. 1C) and then adding the duration of neighboring pairs of closed and open events to the histogram (Fig. 1D). For a given Markov model, rate constants are drawn from within the parameter space and a time series is simulated (*x_sim_*[*t*]). A 2D-histogram is created from *x_sim_* [*t*] analogously to *x_exp_* [*t*] (Fig. 1D,E) and the two histograms are compared using the given objective function (illustrated in Fig. 1F,G). A genetic algorithm is used for optimizing the objective function. The reasoning behind using such a computationally demanding algorithm is that the objective function is considerably affected by the stochastic nature of the simulation process, illustrated in Fig. 1G, with the difference of two dwell-time histograms derived from the same model. Other algorithms, like the Simplex-Algorithm, fail to navigate the complex search space of this task (Huth et al., 2006). In addition, with utilizing a High Performance Computing (HPC) Cluster, analysis of *x_exp_*[*t*] was now conducted with ensemble fits for exploring multiple fit solutions, considering the global optimum and for statistical analysis.

**Figure 1.**
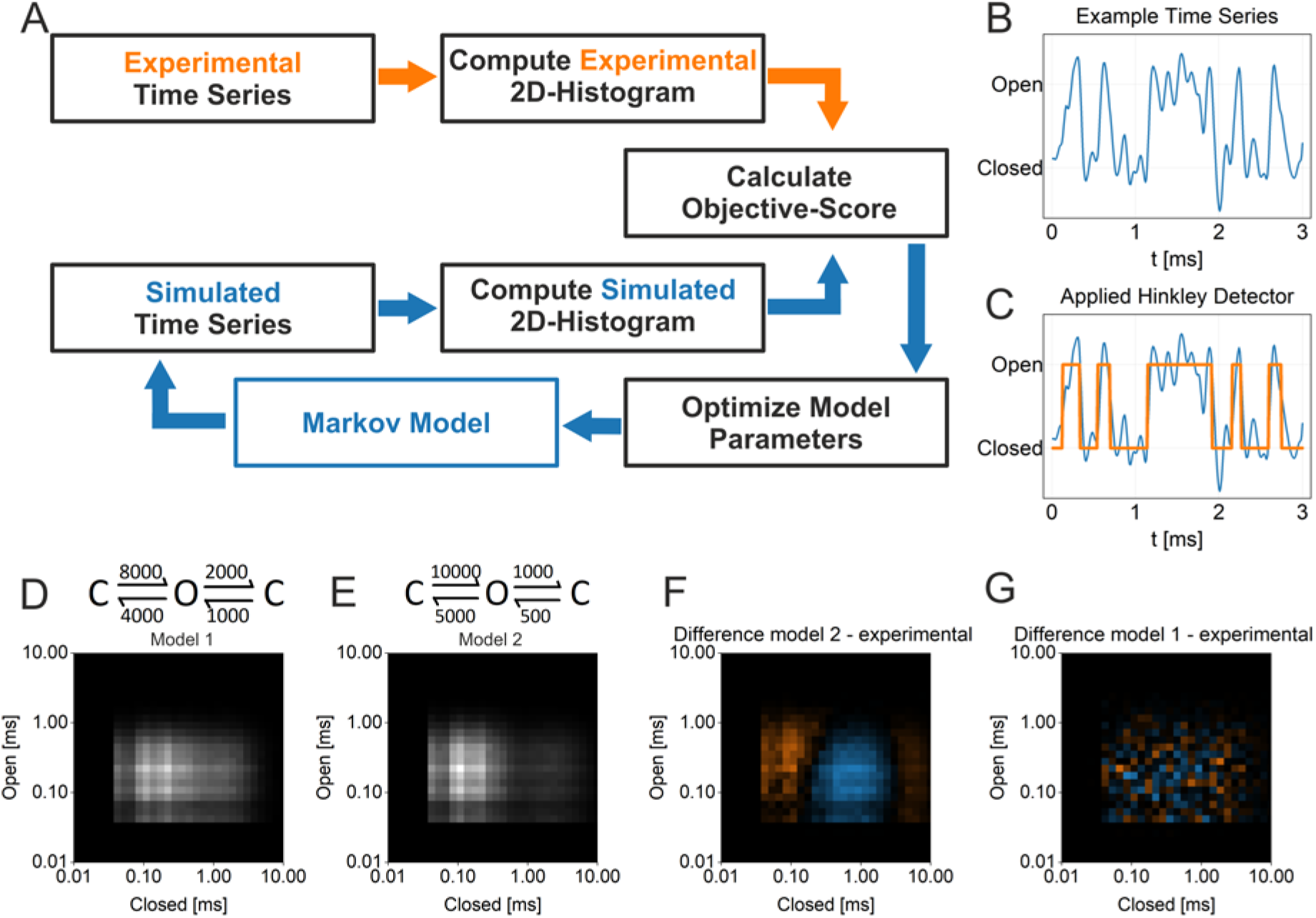
Illustration of the 2D-Fit algorithm. **A** Flowchart of the 2D-Fit. The scheme depicts the iterative simulation method to estimate kinetic parameters from single-channel data as previously proposed (Magleby and Weiss, 1990b) and used in this study. Inputs are an experimental time series *x_exp_* [*t*] and a given Markov model for simulation of time series *x_sim_* [*t*]. The outputs are the optimized model parameters, namely the rates *kij* and if applicable the current of the open state. Color code range is 0 (black) to 2600 (white) **B** Sample of a simulated time series *x_sim_*[*t*] generated with the Markov model 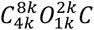 (in Hz) with SNR = 5. Color code: 0 (black) to 2600 (white) **C** Depicts the same section as in B with application of the Hinkley jump detector (Schultze and Draber, 1993) to generate an idealized time series (orange). **D**, **E** Typical 2D-histograms are illustrated, derived from two simulated time series with the Markov models stated above. **F,G** Finally, the application of the objective function (Eq. 3) is illustrated by F subtracting the histograms in (D,E) and by G subtracting histograms from two time series generated with the same model in D. **F** substantial deviation of *x_exp_*[*t*] and *x_sim_*[*t*] color coded from −726 (blue) to +726 (orange), and **G** perfect match showing stochastic deviation only, color coded from −156 (blue) to +156 (orange).

### Simulation of time series

A patch-clamp time series consists of transitions between the closed state and one or more open states with discrete current levels in the order of fA to pA. Transitions between states are governed by conformational changes of ion channel proteins, termed gating. To improve the signal-to-noise ratio a low-pass filter has to be applied considerably limiting time resolution. Time series, both pseudo experimental *x_exp_*[*t*] and simulated *x_sim_*[*t*] were simulated with a 12-bit resolution according to a given Markov model with connected open and closed states and rate constants *k_ij_* governing the transitions between states. In addition, the length of the time series, closed and open channel current levels, noise amplitude, as well as sampling and filter frequency are specified. In the first step, a noise-free time series is generated with discrete current levels as described previously (Schröder et al., 2005). Then, a digital Bessel step response is applied to each transition between current levels. Finally, a fulllength time series consisting of low-pass filtered white noise adjusted to the signal-to-noise ratio (SNR) was superimposed. All time series used in this study were generated with a sampling frequency of 100 kHz and a low-pass filter with the cut-off set to 10 kHz, consisting of ten million samples (100 s) if not stated otherwise. Because of the length, we consider the assumption of microscopic reversibility generally satisfied.

The following changes were made regarding the previous version of the program (Huth et al., 2006). The random generator was updated to a 64-bit Mersenne Twister engine (mt19937). Noise was generated in full length for each simulated time series to prevent systematic artifacts. For the low-pass filtering of noise, a Python 4-pole digital Bessel filter routine (scipy.signal.bessel) was adopted.

The SNR used throughout is defined as

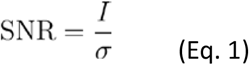

with *I* being the current amplitude between closed and open level and *σ* the standard deviation of the low-pass filtered white noise equaling input noise level for simulation purpose.

### Generation of 2D-histograms

The idealization of the time series was done with a Hinkley jump detector (Schultze and Draber, 1993) yielding discrete current levels. Dwelling in a current level with a lifetime Δ*t* is termed event. The lifetimes of two neighboring events (Δ*t_n_*, Δ*t*_*n*+1_) are combined into pairs of variates. The distribution of variates is then depicted in a logarithmically binned histogram with the axes corresponding to the dwell-times of closed and open events. (Labarca et al., 1985; Magleby and Weiss, 1990a). Since microscopic reversibility is assumed for the time series both closed-open and open-closed pairs are included in the same histogram. A resolution of ten bins per decade has proven to be adequate for the histograms (data not shown) yielding typically a 50 x 50 matrix (from 1 μs to 100 ms). Importantly, the procedure for generating the histograms is exactly the same for the experimental as well as the simulated time series *x_exp_*[*t*] and *x_sim_*[*i*]. The objective-score, calculating the correlation of histograms generated with *x_exp_*[*t*] and *x_sim_*[*t*] was finally calculated according to either the LogLikelihood (LLH)

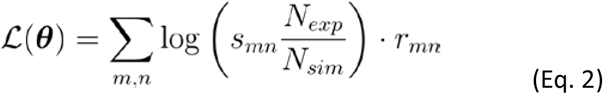

or the 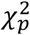

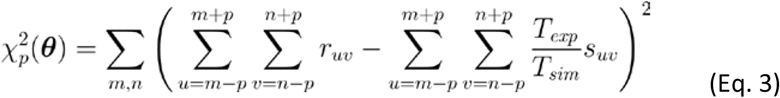

with *r_uv_* and *s_uv_* the bin occupancy in the experimental and simulated 2D-histogram respectively for column *u* and row *v, N_exp_* and *N_sim_* the total number of events, *T_exp_* and *T_sim_* the length of the time series in the experimental (exp) and simulated (sim) time series or histogram respectively, ***θ*** the vector comprising all model parameters, i.e. the rates *k_ij_* For simplicity, boundary conditions (such as padding) were not included in (Eq.3). We also used the mean absolute percentage error (MAPE), as a fit-independent metric for comparing and evaluating the precision of rate constants estimates. It is defined as

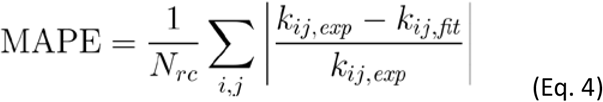

with *k_ij,exp_* the rate constants of the given pseudo-experimental time series, *k_ij,fit_* the rate constants determined by the fit algorithm, the state indices *i, j* and *N_rc_* the number of rate constants of the given model. Only valid transitions are considered in the sum, i.e. combinations of *i, j* for which transitions in the given topology exist (*k_ij_* ≠ 0 and *i* ≠ *j*)

### Genetic algorithm

A genetic algorithm was used to optimize the objective function. The genetic algorithm library Galib (Wall, 1996) was employed with the settings as indicated in table 1.

**Table 1.** Settings of the genetic fit algorithm related to the Galib library (Wall, 1996). If not otherwise stated settings from the table were used. Parameter denoted by “*” or “**” were optimized according to a serialized optimization process (“*” data not shown).

| 2D-fit algorithm | Simple Genetic Algorithm |
| --- | --- |
| Population size | 800 |
| **Objective function | LLH (Eq. 1) |
| **Stopping criterion | 40 generations without improvement |
| **Crossover rate | 0.600 |
| **Mutation rate | 0.010 |
| *Elitism setting | True |
| *Scaling of rate constants | Uniform logarithmic |
| *Parameter coding | GABin2Dec, 16 Bit |
| *Distribution of initial parameter space | Uniform |
| *Selection function | No scaling, Tournament Selector |
| *Crossover | EvenOdd |
| *Mutator | Flip Mutator |
| Oversampling | None (*length of simulated time series/measured time series = 1*) |

### Current level fit of the conducting state

The fit of the conducting (open) state current level was implemented as follows. In addition to the rate constants, the current level is included in the genome of the genetic algorithm. Note the setting of the level does only affect the simulation of time series and not the jump detector where the level is initially fixed close to the anticipated level. We tested different settings of the level for the jump detector at 20% off the actual level in both directions for a 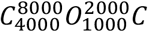 (in Hz) model and did not find significant alterations in fit results (data not shown).

### Amplitude fit

This option enables the calculation of the difference between the current distributions of *x_exp_*[*t*] and *x_sim_*[*t*] (amplitude-fit) according to

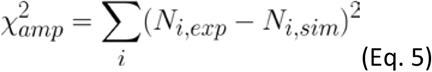

with *N_i,exp_* and *N_i,sim_* being the occupancy of bin *i* ∈ {0,1,2,…,4096} (12 Bit) in the histogram of the experimental and simulated time series respectively. During the fitting process, the resulting deviation (Eq.5) can either be used stand-alone as an objective function or combined with the 2D-histogram objective functions (Eqs. 2, 3). When combined with the LLH (Eq. 2), the LLH is divided by the 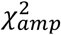. For combination with 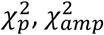 and 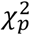 are multiplied.

### MPI implementation

We implemented a Message Passing Interface (MPI) with a manager process governing the simple genetic algorithm and sending out parallel tasks to a variable number of worker processes. The workers are initialized with the experimental time series *x_exp_*[*t*], all settings related to the simulation of the time series *x_sim_* [*t*] including the Markov model, and the selected objective function. Then, they receive the rate constants of the current generation from the genetic fit algorithm and if enabled the current level of the conducting state (level fit). Time series are simulated, 2D-histograms of *x_exp_*[*t*] and *x_sim_* [*t*] are constructed and the objective score according to the objective function is computed. Finally, the objective score is sent back to the manager process. After receiving all results from the current generation, the manager evaluates the objectives scores, generates the next generation and again sends out the new tasks. When the convergence criterion is reached, the manager sends out a number of tasks with the final solution to obtain the mean of the result. Note, with this implementation the reasonable number of parallel worker processes is limited by population size of the genetic algorithm. If not otherwise stated the number of parallel processes was below 50% of the population size.

### Simulations on High Performance Computing (HPC) Cluster and ensemble runs

For ensemble runs, we utilized either the Emmy or the Meggie parallel cluster of Regionales Rechenzentrum Erlangen, Germany (RRZE) with one up to a maximum of 64 nodes. For the Emmy cluster, each node consists of two XEON 2660v2 “Ivy Bridge” processors (Intel) with 10 cores per chip running at 2.2 GHz with a maximum of 40 simultaneous processes if utilizing hyper-threading. For the Meggie cluster each node consists of two Intel Xeon E5-2630v4 “Broadwell” chips with 10 cores per chip running at 2.2 GHz. Hyper-threading was not enabled on the Meggie Cluster.

The C/C++ code of the former program Kielpatch (Huth et al., 2006) was modified and updated to compile within a modern 64-bit LINUX environment using the GCC 8.1.0 compiler (The GNU Compiler Collection) and Open MPI 3.1.1. (Open Source High Performance Computing). The code of the actual program 2DFit64 was compiled using the flag -O2. For batch operation on Meggie, we used the Torque scheduler (Adaptive Computing), and on Meggie Slurm (SchedMD) workload manager. Typically, an ensemble fit with 50 runs completed within 24 h using eight nodes (320 processes, 800 population size, 10 million samples per time series). Note, the computational demand increases steeply with higher rate constants.

### Data analysis

Data analysis was performed using Origin PRO2021b software (OriginLab Corp). All data are expressed as mean ± standard error of the mean (SEM) if not otherwise stated.

## Results

### Estimating parallel performance

Modeling patch-clamp data with the 2D-fit (Fig. 1) is accomplished by simulation of time series and adjusting the rates of the given Markov model until experimental and simulated time series are matching (Fig 1). Due to the stochastic nature of the simulation, the objective function used for evaluation of derived 2D-histograms is endowed with a considerable variation given the same set of rate constants. To cope with this challenge, we employed a genetic algorithm for fitting purpose (Wall, 1996) resulting in a sizable amount of time series to simulate. In this study, we distributed time series simulations to many parallel processes using a Message-Passing Interface (MPI). The implementation of a simple genetic algorithm (Wall, 1996) means in terms of parallel simulation that one generation has to be fully evaluated before commencing with the next generation. Therefore, the population size sets the upper limit for the number of parallel processes. The first task was to evaluate parallel capabilities of the 2DFit64 program. Performance was analyzed by utilizing an increasing number of parallel nodes while keeping the workload per process constant. Overall, computational load was scaled by increasing population size. Ideally, with this task, computation time should stay the same. As outlined in Fig. 2 the drop in parallel performance utilizing more and more processes was insignificant and below 5% up to a total of 64 nodes tested. Note, the increase in computation time manifested mainly in the first generation indicating that it was due to initial loading of required files. As expected, the more recent architecture of the Meggie cluster performed slightly better and showed less drop in parallel performance. Overall, scaling of the fit by increasing population size in parallel with the number of nodes comes without a major penalty.

**Figure 2.**
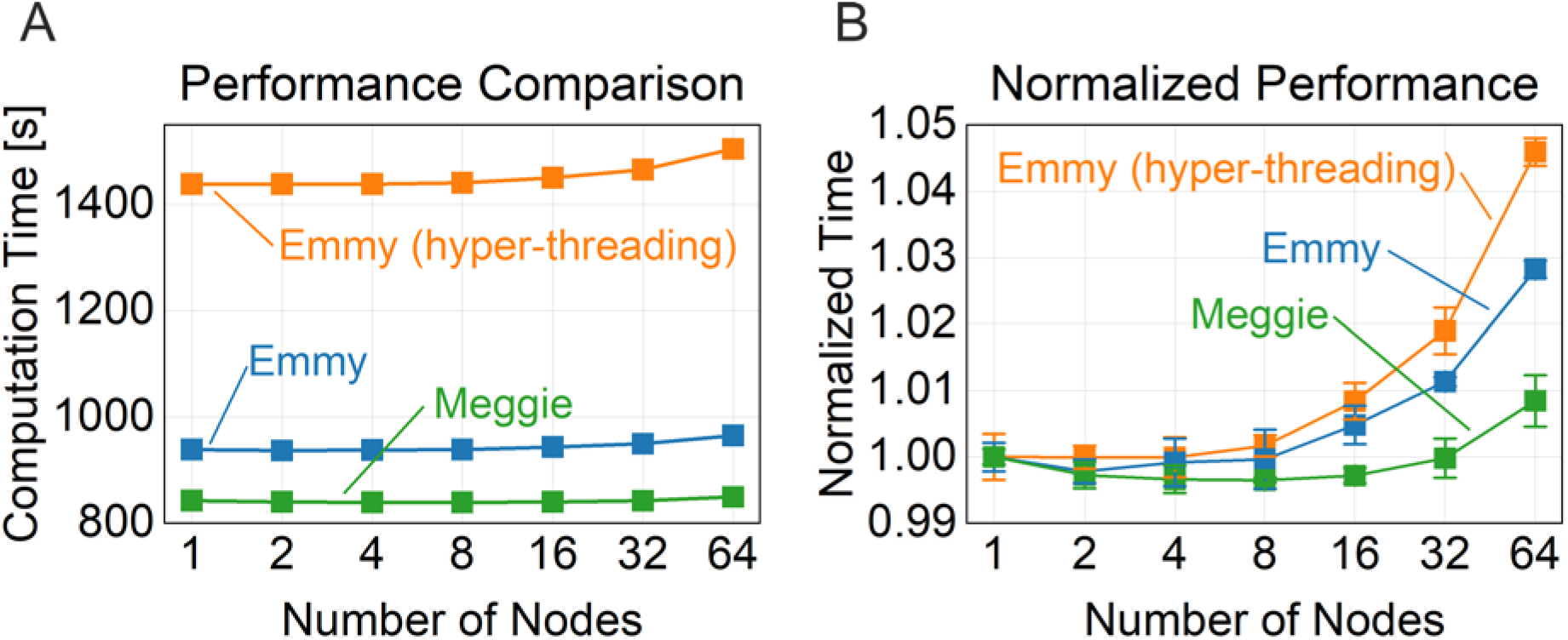
Fit performance using HPC-Cluster Emmy and Meggie drops marginally by utilizing an increased number of nodes. The number of nodes from one up to 64 was increased while the workload per node was kept constant by scaling the population size of the genetic algorithm. Accordingly, the number of processes ranged from 20 to 1280 or from 40 to 2560 with hyper-threading. Notably, the differences in parallel performance were mainly due to initial loading of required files. **A** The total computation time is depicted for the Emmy cluster with and without hyper-threading and for the Meggie cluster without hyper-threading. **B** Total time was normalized to the mean for one node. For this task a 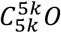 (in Hz) model was used with the time series containing 10M samples. The number of experiments was n = 4 and each data point is depicted as mean ± SD.

### Ensemble fits and objective functions

The next section is related to different objective functions operating on 2D-histograms and additionally, a measure of fit-precision independent of these functions. The previous versions of the 2D-fit (Magleby and Weiss, 1990b; Huth et al., 2006) were severely limited by the lack of computing power in a fundamental way. Single solutions obtained were not reliable due to the stochastic nature of the simulation process and the resulting variable outcome of the fitting algorithm. This issue was now overcome by using ensemble fits. This allowed us to compare different scenarios by statistical means and, importantly, to identify possible global solutions with high confidence. In the first step, we optimized the basic settings of the genetic algorithm in a serial procedure that were then kept for all subsequent fits (table 1). We then conducted a large ensemble of 1000 individual fits of a known time series generated with a relatively complex five-state Markov model. The fits were ranked according to the objective function, in this case the LogLikelihood (LLH, Eq. 2, Fig 3A). The horizontal orange line indicates the LLH obtained with values of the given model denoting the highest possible LLH. The prototypical continuous and shallow curve of the fit data (Fig. 3A), reaching the horizontal line lets us reasonably assume having found several solutions very close to the global maximum. Note, the pseudo experimental time series used for fitting is also subject to statistical variations that are not noticeable at this scale. With the ensemble, it is now clear, that only part of the solutions were close to the maximum. This clearly emphasizes the necessity of ensemble fits. In parallel, we computed the mean absolute percentage error on the obtained rates (MAPE, Eq. 4) for the same set of data as independent measure of fit accuracy (Fig. 3B). The MAPE correlates well with the objective score albeit considerable fluctuations, at least in the range of the best solutions with lowest ranks. Again, the MAPE indicates that within the ensemble roughly 20 percent of the solutions are within 10 percent of deviation that we consider very good for this task.

**Figure 3.**
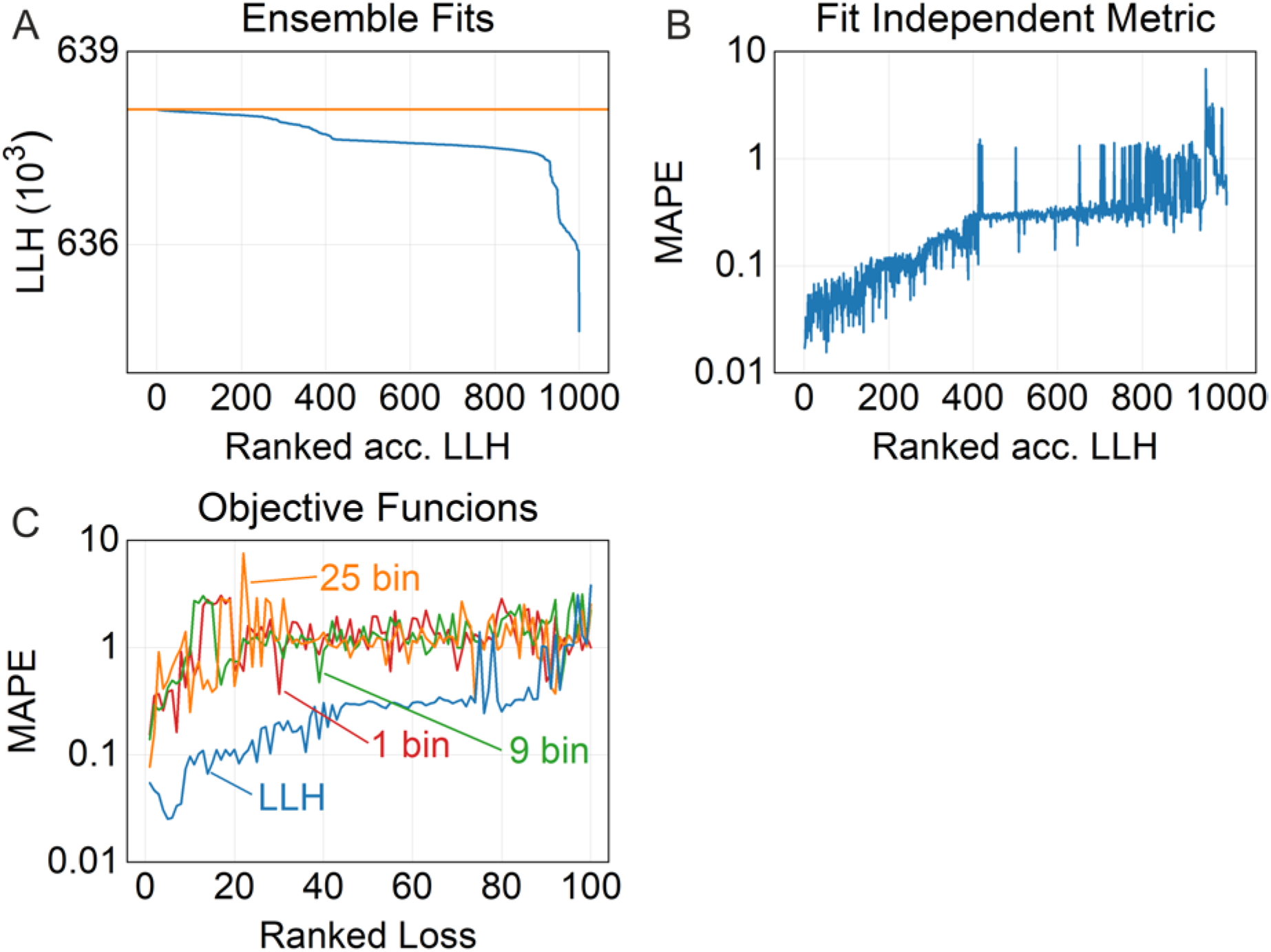
Evaluation of fit performance using ensemble fits and the fit-independent metric mean absolute percentage error (MAPE). A pseudo experimental time series *x_exp_* [*t*] was generated using the five state Markov model 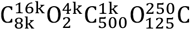 (in Hz) with 10M samples and SNR = 5. The boundaries for rate constant estimation were *k_ij,min_* = 10 *Hz* and *k_ij,max_* =100 *kHz* the settings of the genetic algorithm were according to table 1. **A** Fit of the experimental time series with an ensemble size of 1000 repeats using the LLH objective function (Eq. 2). The results were ranked according to the objective score. The orange horizontal line indicates the LLH for rates as given for the model of the pseudo experimental time series (mean of 10000 repeats, SD is too small for visualization). **B** The fit-independent score MAPE (Eq. 4) was computed from the data in A and displayed with identical ranking. **C** The same time series as investigated in A was fitted using LLH or 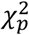 objective functions (Eq. 2, Eq. 3 with *p* ∈ {1,2,3} corresponding to an average filter of 1, 9 or 25 bins) with an ensemble size of 100. The data was ranked according to the respective objective score. The MAPE (Eq. 4) was computed for comparison.

In our previous study we used a filtered 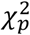 criterion (Eq. 3) for the objective function in contrast to Magleby and Weiss (1990b, 1990a) using the LLH (Eq. 2). The reason was that close to the global solution 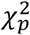 could better discriminate compared to the LLH (Huth et al., 2006). In this study, we compared the different objective functions for a given fit problem using the MAPE for evaluation. Ensemble fits revealed that the LLH overall yielded better solutions compared to 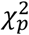 (Fig. 3C). A possible explanation could be that away from the global solution the error scape for the 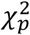 criterion might be rather shallow.

### Performance of the genetic algorithm

In this section, we consider the settings of the genetic algorithm that likely have the biggest impact on the fit performance, population size, stopping criterion, crossover rate, and mutation rate. We used the same time series of a five-state model from the previous section, all other settings from table 1, LLH as objective function (Eq. 2), and ensembles of 100 fits. Fig. 4A indicates that with a comparable small population size of 200, still some good solutions could be obtained. As expected, increasing population size increased the number of good solutions while decreasing the number of outliers. We note here that the mean number of generations before reaching the stopping criterion increased moderately with larger population size indicating more evolution taking place (from 178 at 200 population size to 192 at 3200 population size). Overall, increasing population size is very costly in terms of computation. The main message is that small population size requires larger ensemble fits and vice versa. From the results in Fig. 4A we consider a population size of 400 and an ensemble size of 100 as feasible for this model. However, for distinct complexity of the model the best compromise between both settings might be different.

**Figure 4.**
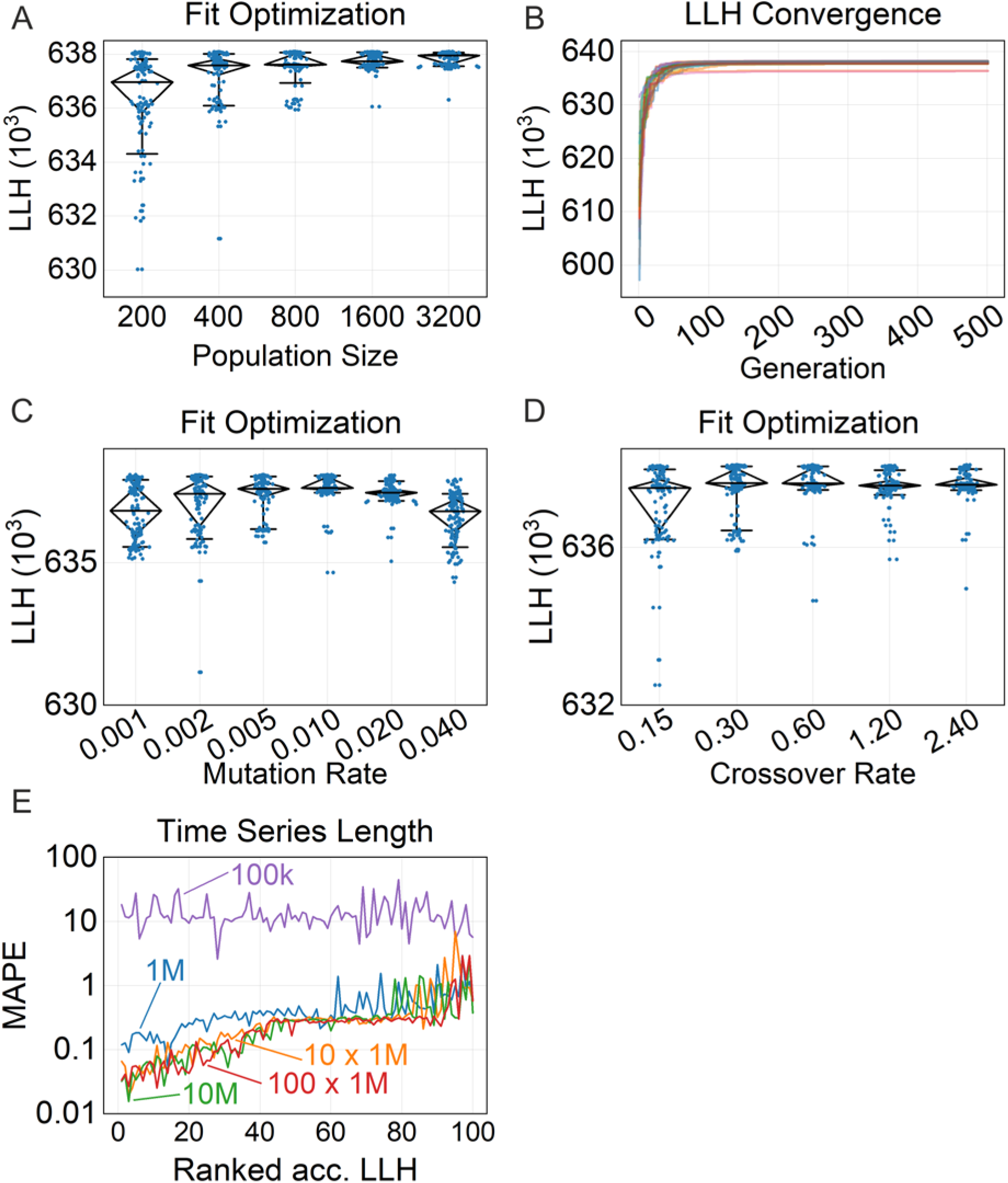
A-D Optimization of the cardinal settings of the genetic algorithm. A pseudo experimental time series *x_exp_* [*t*] was generated using the five state Markov model 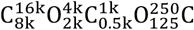 (in Hz) with 10M samples and SNR = 5. The boundaries for rate constant estimation were *k_ij,min_* = 10 *Hz* and *k_ij,max_* = 100 *kHz*. The fit ensemble consisted of 100 repeats **A,C,D** and 25 repeats **B**. The LLH objective function (Eq. 2) is depicted using different population size **A**, crossover rate **C**, and mutation rate **D**. All other settings were according to table 1, respectively. B shows the LLH loss over 500 fit generations without the stopping criterion enabled. **E** Estimating the length of the experimental time series required for obtaining meaningful fit results. The Markov model and all other settings were taken from the experiments in (**A-D**). Ensemble fits of 100 repeats were conducted for time series of 100k, 1M and 10 M samples. In addition, we tested over-sampling of the simulated time series *x_sim_* [*t*] simulated 10 or 100 times longer than the experimental time series *x_exp_* [*t*] of 1M samples.

The stopping criterion is implemented in such a way that the genetic algorithm stops after a given number of generations without improvements. Partially, improvements were due to finding a better objective score after reevaluating an existing solution within the genome due to stochastic variability. Using the same model as before Fig. 4B depicts an ensemble of 25 fits, proceeding for 500 generations without the stopping criterion enabled. Most of the fits converged quickly with constant improvement of the objective score until about 100 generations. Thereafter hardly any improvements could be observed. We conclude that a stopping criterion of 40 generations is a rather conservative setting for this model to reach the plateau.

Crossover (“mating”) and mutation are two fundamental mechanisms altering the genome, to yield potentially better solutions in the next generation (Rechenberg, 1978; Holland, 1992). We assume that an optimum exists for both parameters. With ensemble fits, we tested different settings using the five-state Markov model from the previous tasks. Within the tested range the LLH showed a rather weak dependence on both parameters (Fig. 4C,D). However, the ensemble fits indicate the same settings as estimated in the initial optimization process. With help of the ensemble fits, the settings in table 1 represent a close to optimal tuning of the algorithm.

### Length of time series

Next, we determined how long an experimental patch-clamp time series must be for successful modeling. In addition, we tested the option to use oversampling, meaning *x_sim_* [*t*] was simulated n- times longer than *x_exp_*[*t*]. Using HPC resources is now a feasible option. Oversampling reduces the relative statistical variation in the simulated 2D-histograms.

For a time series of 100k samples corresponding to 1s recording time and 1269 events for this example we were not able to obtain meaningful solutions indicated by the MAPE (Fig. 4E, purple trace). Increasing the number of samples 10-fold (12817 events, blue trace) allowed the fit to converge to proper solutions. Using a 10-fold oversampling (orange trace) or increasing the length again 10-fold (green trace) provided solutions with better accuracy. However, 100-fold oversampling of 1M samples (red trace) did not further decrease the MAPE. In summary, we can now estimate the minimal length of the time series required and provide a method to significantly improve modeling results using moderate oversampling.

### Estimating fast gating events

Because of the tiny currents in the order of fA to pA, noise is a fundamental issue of single-channel recordings (Parzefall et al., 1998). To keep noise levels at reasonable range, low-pass filtering has to be employed resulting in a reduction of effective bandwidth. If events are shorter than the rise time of the filter (fast gating) they are distorted resulting in an apparent reduction of the current amplitude. The jump detector therefore might miss events. On the other hand, a high noise amplitude leads to false alarms. The setting of the low-pass filter is therefore always a compromise. In previous work it was shown that potentially errors related to the jump detector cancel out to a certain degree for *x_exp_*[*t*] and *x_sim_*[*t*] making the 2D-fit robust in this regard (Magleby and Weiss, 1990b; Huth et al., 2006). In addition, rates considerably beyond the filter frequency could be estimated. Here we want to explore the capabilities of the 2D-fit to extract fast rates beyond the filter frequency using a COC three state Markov model, expanding the two-state topology of the model used in the previous study (Huth et al., 2006). Importantly this approach required the extraction of fast and slow rates in parallel therefore adding considerable complexity. Fig. 5A illustrates a time series generated with rates below the limit of the low-pass filter and Fig. 5B with rates significantly beyond the limit resulting in an apparent current reduction. Ensemble fitting was conducted on time series with increasing rates *k_12_* and *k_21_* connecting C_1_ and the O_2_ state while keeping the other rates constant (slow rates). The mean LLH was slightly increased for the ensembles with higher rates (Fig. 5C) and then decreased with a very sharp drop for the fastest pair of rates (128 kHz, 256 kHz). The calculated MAPE was mostly inverse to the LLH (Fig. 5D). With these results we have now established the limit of the 2D-fit to extract rates beyond the cut-off frequency of the low-pass filter which is around 10-fold. We then artificially reduced the current amplitude of the jump detector and fitted the time series again. First off, with the reduced level, we approximately matched the apparent current reduction of the fastest rates and secondly, we wanted to know whether exact current level is a prerequisite of the algorithm to function properly. As stated in the methods section, setting the jump detector 20 percent off the simulated current level did not affect fit results. However, the previous experiment was conducted with rates below the cut-off frequency. Our data indicated that with the reduced level of the jump detector we still achieved meaningful results, albeit with reduced LLH (Fig. 5E) and reduced accuracy (Fig. 5F).

**Figure 5.**
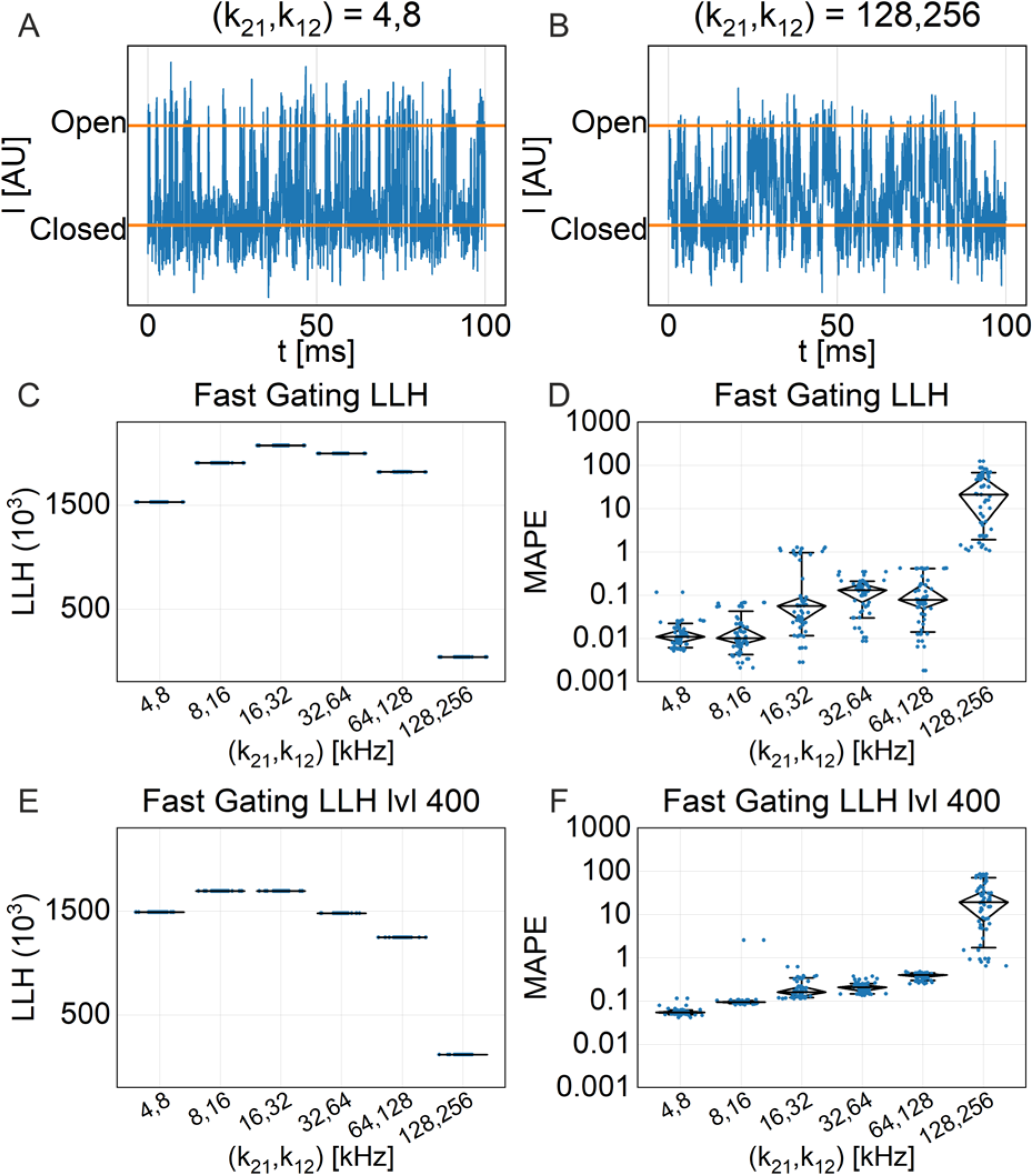
Capabilities of the 2D-fit to determine fast rates. Time series were generated with a 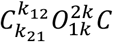 (in Hz) model with a length of 10M samples, the current amplitude of the open state set to 500 AU, and SNR = 5. *k_12_* and *k_21_* were set as stated. The other parameters of the 2D-fit were according to table 1, with population size = 800 and with an ensemble size of 50. **A,B** Example time series sections with (*k_12_*, *k_21_*) = (4, 8) and (128, 256), respectively. The closed and open current levels of the simulation are indicated by the orange horizontal bars. Note the apparent reduction of the open state in **B**. **C** depicts the LLH objective score of the ensemble fits depending on *k12* and *k_21_* as indicated and **D** the calculated MAPE. **E, F** Distinct from the previous experiments the level detector of the 2D-fit program was arbitrary set to 400 AU instead of 500 AU with **E** depicting the objective score (Eq. 2) and D the calculated MAPE (Eq. 4).

### Sensitivity to noise

We explored the performance related to the SNR in more detail with ensemble fits using two Markov models and varying noise levels. The first set of time series *x*_exp *i*_ [*t*] was generated with a 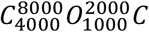 model and different SNR ranging from 0.25 up to 8. All rate constants were kept below the frequency of the low-pass filter (illustrated in Fig. 6A-C). The second set was generated with the same topology and the same SNR values, but two rate constants were increased markedly beyond the filter frequency (fast gating, illustrated in Fig. 6D-F). As expected, the LLH increased considerably with increasing SNR (Fig. 6G,I). For comparing the fit performance at different SNR, we applied the MAPE (Fig. 6H,J). As a reference, we included the MAPE of randomly drawn logarithmically distributed rate constants from within the parameter space. To obtain meaningful results a SNR of at least 1 was required with the accuracy improving steeply with increasing SNR. The task with fast gating had the same dependency on SNR except for the estimated rates being less accurate overall. The results imply that fast rates can still be extracted from time series with relatively high noise.

**Figure 6.**
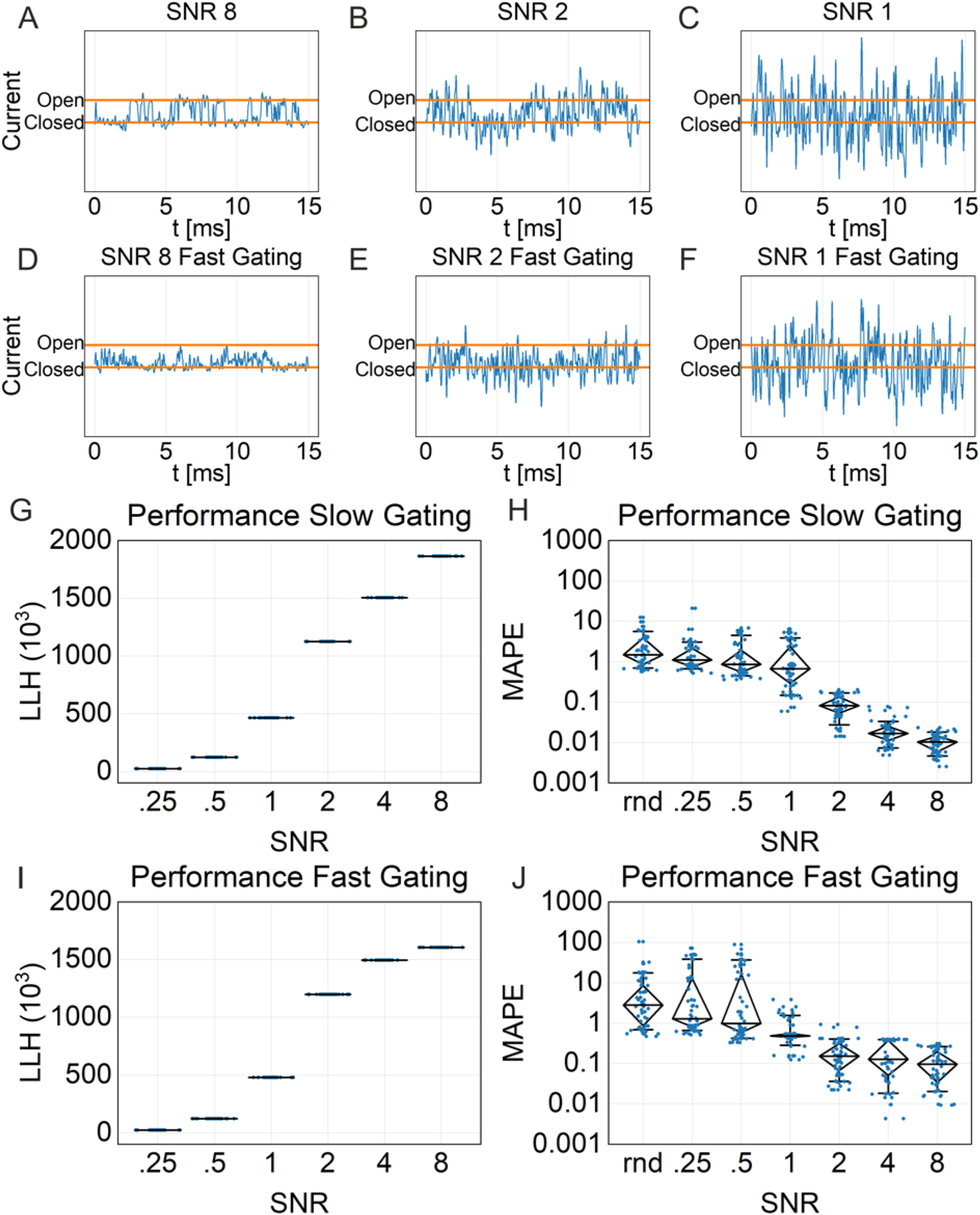
Performance of the 2D-algorithm related to noise and in combination with fast gating. Pseudo experimental time series *x_exp_* [*t*] from a COC Markov model with different SNR as indicated were generated. The time series contained 10M samples. For investigating slow gating processes, the model was 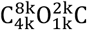 (in Hz) and for fast gating 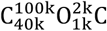 (in Hz). **A-F** Illustration of pseudo experimental time series with the SNR as indicated. **A-C** slow gating, D-F fast gating. **G-J** *x_exp_* [*t*] were analysed with the 2D-fit with an ensemble size of 100. The fit boundaries for all *k_ij_* were 10 Hz to 100 kHz for the slow gating model and 100 Hz to 1 MHz for the fast gating model. All other settings were according to table 1. **G,I** The LLH (Eq. 2) was estimated for fit ensembles with different SNR (Eq. 1) as stated in the graphs. **H,J** From the data in **G,I** the MAPE (Eq. 4) was computed and plotted against the SNR. As reference, we included the MAPE for randomly drawn rates *k_ij_* from an exponential distribution. Diamond charts indicate the median, 25 and 75 percentile and whiskers the 10 and 90 percentiles.

### Automated level detection and amplitude fit

Ion channels operate on a time scale, which, in part, cannot be fully resolved in patch-clamp recordings. Nevertheless, fast gating processes close to the time scale of molecular dynamic simulations are of great interest. At least two methods are capable of extracting rate constants far beyond the frequency of the low-pass filter - the 2D-fit used here (Huth et al., 2006) and the fit of β- distributions (Schröder et al., 2005; Schroeder, 2015). Fast gating events cause a deviation from the Gaussian distribution of current amplitude histograms. The skewness of the β-distribution can be explored to deduce fast rate constants. However, both algorithms require exact knowledge of current levels. While the closed state can generally be inferred from episodes without gating, the current level of the open state might not be accessible due to an apparent current reduction caused by low-pass filtering, as mentioned above. In our previous study we have outlined an iterative approach with the 2D-fit (Huth et al., 2006) and applied it to gating processes of voltage gated sodium channels (Huth et al., 2008). This approach is very time consuming and might not be feasible for all recordings. In the previous section we demonstrated that the objective function is potentially sensitive to the correct setting of the level for the jump detector in case of fast gating (Fig. 5C vs. 5E). Therefore, we have advanced the 2D-fit with a new capability. The current level of the conducting state can now be included in the genome of the genetic fit. It was implemented in such a way that it only affects the simulation of time series, not the jump detector. The level is then fitted, within given boundaries, simultaneously with the rate constants *k_ij_*. Secondly, the current distributions of *x_exp_*[*t*] and *x_sim_*[*t*] were calculated (current amplitude histograms, CAH) and then compared according to least squares deviation (Eq. 5). The CAH deviation can be used as a stand-alone objective function, which is analogous to fitting β-distributions, or combined with 2D-histogram objective functions (see methods). For testing the capabilities of these new features, we analyzed time series generated from one model with slow rates below the cut-off frequency of the low pass filter (Fig. 7) and another with fast rates causing an apparent current reduction (Fig. 8). In addition, for the model with the slow gating process we tested two different SNR as illustrated in Fig. 7A,D.

**Figure 7.**
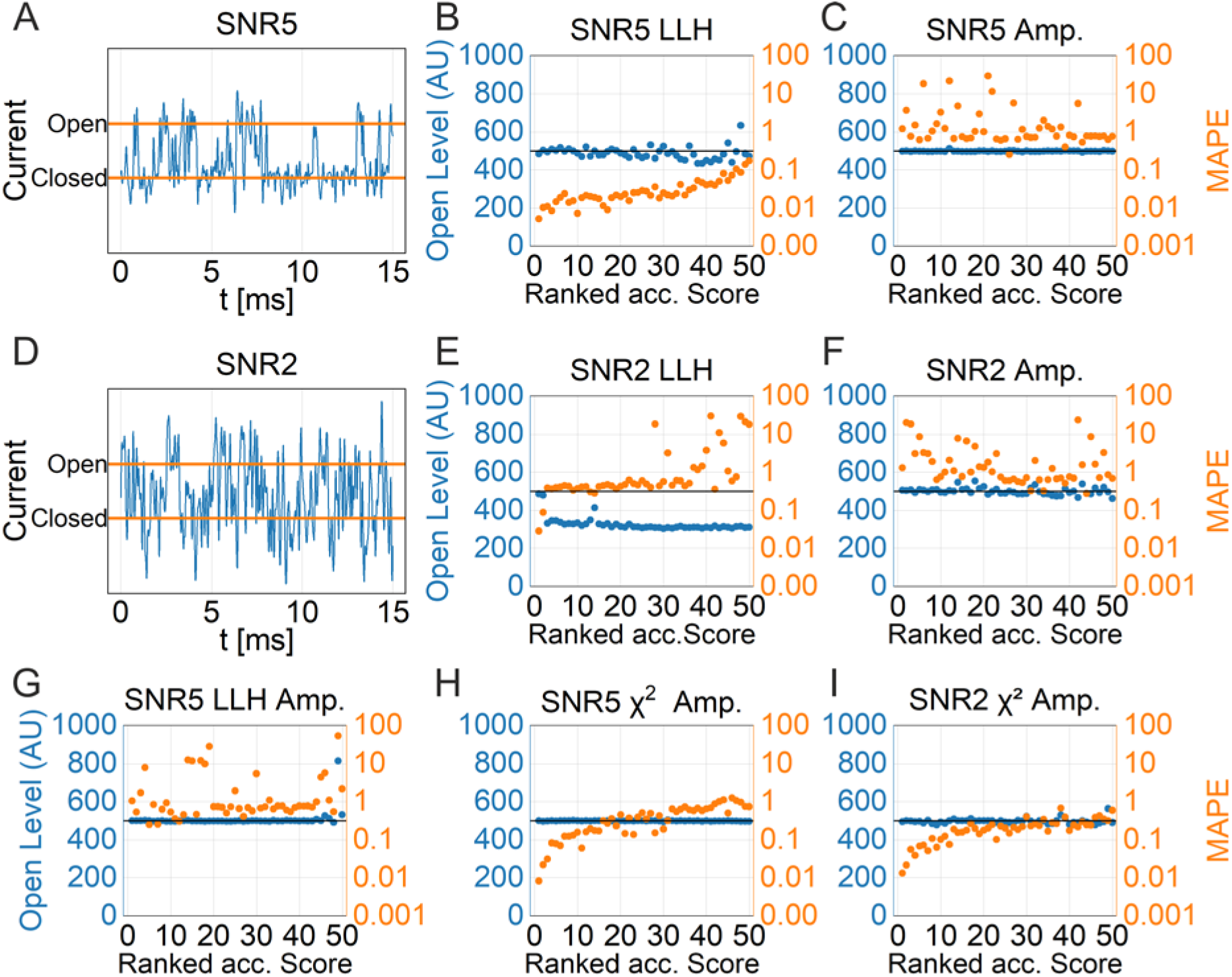
Current level estimation with an underlying slow gating process. Two pseudo experimental time series *x_exp_*[*t*] were generated with the Markov model 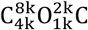 (in Hz) using a different signal to noise ratio (SNR). The length of the time series was 10M samples. The current amplitude of the open state was 500 AU. Rates *k_ij_* in addition to the current level were estimated with the 2D-fit with an ensemble size of 50 using different calculations of the objective function. The fit boundaries for all were 1 Hz to 1 MHz and the boundaries for level estimation ranged from 1 AU to 1000 AU. All other settings were according to table 1. **A,D** depict a short period of the time series with **A** SNR = 5 and **D** SNR = 2. The orange bars indicate closed and open current levels, respectively. **B,C,E-I** The different objective functions used in the 2D-fit are stated for each graph. MAPE (orange) and estimated level (blue) are plotted against the ranked objective score. LLH according to Eq. 2, χ^2^ according to Eq. 3, Amp. is the difference between the current histograms *x_exp_* [*t*] and *x_sim_*[*t*] according to Eq. 5. If combined, LLH was divided by the Amp. score whereas χ^2^ was multiplied with the score.

**Figure 8.**
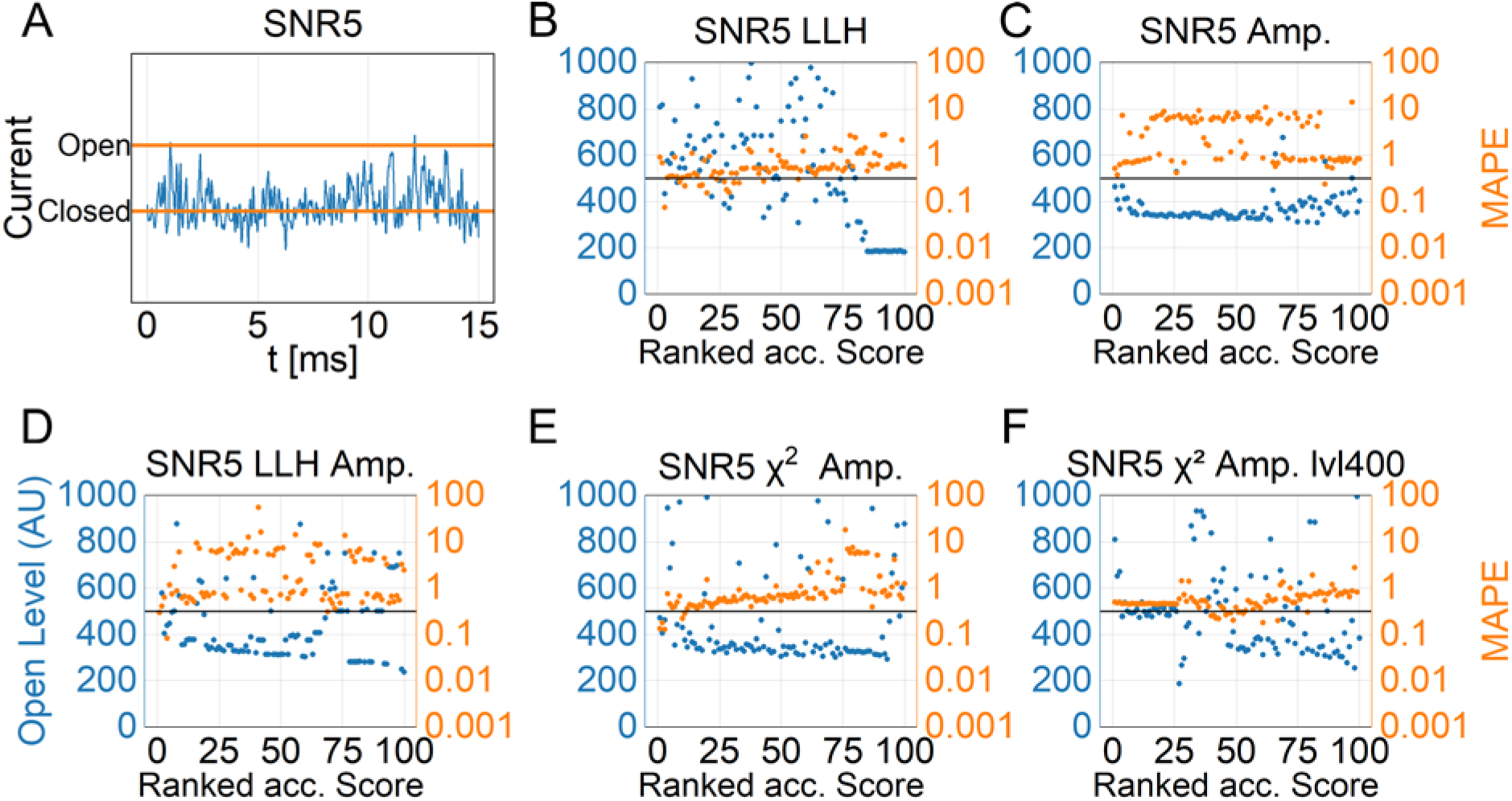
Current level estimation with an underlying fast gating process. A pseudo experimental time series *x_exp_* [*t*] was generated with the Markov model 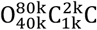 (in Hz) using a signal to noise ratio SNR = 5. The length of the time series was 10M samples. The current amplitude of the open state was 500 AU. Rates *k_ij_* in addition to the current level were estimated with the 2D-fit with an ensemble size of 100 using different objective functions. The fit boundaries for all *k_ij_* were 1 Hz to 1 MHz and the boundaries for level estimation ranged from 1 AU to 1000 AU. All other settings were according to table 1. **A** depicts a short period of the time series. The orange lines indicate the closed and the open states. **B-F** MAPE (orange) and estimated level (blue) are plotted against the ranked fit-score with the objective function indicated above the graphs. LLH according to Eq. 2, χ^2^ according to Eq. 3, Amp. is the difference between the current histograms *x_exp_* [*t*] and *x_sim_*[*t*] according to Eq. 5. If combined, LLH was divided by the Amp. score whereas χ^2^ was multiplied with the score.

Overall, this task was more difficult compared to estimating rates only, indicated by a reduced number of reasonable solutions in the fit ensembles. For the slow gating process, with an excellent SNR = 5, the 2D-Fit with standard settings, using the LLH objective function was able to detect the current level and the rates (Fig. 7B). However, with a reduced SNR = 2 the 2D-Fit was mostly failing (Fig. 7E). Using the CAH deviation (Eq. 5), fit results always provided good solutions for level detection, irrespectively of SNR (Fig. 7C,F). However, as expected for a slow gating process without deviations from a Gaussian distribution, the rates could not be determined. We then used combined objective functions. The LLH together with the CAH deviation did not improve rate estimation (Fig. 7G). Surprisingly, the combination of CAH deviation and 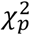 objective function provided the best results for current level and rates estimation (Fig. 7H,I) whereas using the 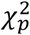 objective function alone did not provide good estimates (data not shown). The reason for the difference in LLH and 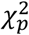 objective-function performance in this task might be different time series normalization (number of events vs. simulated time).

With an underlying fast gating process, current levels where apparently reduced due to low-pass filtering (Fig. 8A). Here, we only tested SNR = 5. Now, without using the current-amplitude-histogram deviation score, level detection failed completely (Fig. 8B). Otherwise with the amplitude fit (Fig. 8C), only the best solutions of the ranked fit ensemble yielded a level estimation close to the original level. Presumably, the combination of slow and fast rates prevented reliable detection of the level and also failed to estimate rates. The same was true for using LLH together with CAH deviation (Fig. 8D). The combination of CAH deviation and 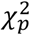 objective functions, which worked best for slow rates, performed in a way that both rates and level could be approximated from the very best solutions (Fig. 8E).

As outlined in Fig. 5F we finally reduced the level for the jump detector. Interestingly enough, the best-ranked solution did not provide good level or rates estimates (Fig. 8E). However, the following solutions were all very similar with a correct level. Level detection for a fast-gating process appears to be very difficult, but not impossible given a good SNR and fit ensembles to provide a basis for selecting proper results. Alternatively, the level estimation taken from Fig. 8E,F could be used in a two-step process, to fit the rates after the level was determined.

### Topology estimation of the underlying Markov model

The likely most difficult task of modeling single-channel time series is addressed in this section. After determining the underlying Markov model of a time series, states and rates can be linked to conformational changes in the ion-channel protein. Therefore, identifying the sequence of states in a Markov model is of evident importance. Typically, conventional 1D-Dwell-Time diagrams allow identifying the number of open and closed states by determining time constants (Qin, 2007). The sequence of states can then be guessed by assumptions of protein structure or by inference from (ambiguous) whole-cell recordings (Lampert and Korngreen, 2014). The situation was improved by introducing 2D-histogramms (Magleby and Weiss, 1990a). Combining open- and closed-dwell times preserves the sequence of states for analytical tools. Still an exhaustive search over all model topologies is required and, in the end, there was no statistical mean to estimate the quality of the obtained solution. In this study, we have now overcome this latter issue by using the 2D-fit together with ensemble fits as described above. Two time series *x_exp_*[*t*] with a four state COCO Markov model were simulated with rate constants as stated in the caption of Fig. 9. In one time series, the rates were kept below the low-pass filter frequency and in the other two rates were increased beyond the filter frequency (fast gating). The second time series aimed to investigate whether fast gating interferes with topology estimation. We performed an analysis over all meaningful linear four state topologies and in addition a COC, a cyclic COCO, and a five state COCOC model. The cyclic and the five state model comprise more rates than the given model and are therefore over-defined. Fig. 9A depicts the results of the topology estimation for the model with slow rates. The ensemble was ranked according to the LLH estimation. To choose the appropriate model, we considered only the best solutions within the ranked ensemble. According to Fig. 9A four models had similar high objective-scores including the correct model COCO. The zoomed in view (Fig. 9B) revealed that among these models the OCCO model had a smaller LLH than the other models. This result was exactly what we expected. The 2D-fit objective score allowed us to discern the correct COCO model along with two other models, cyclic COCO and COCOC that provide (at least) the same degree of freedom. We note here, in this setting, over-fitting does only provide equal but not better LLH. In addition, models with more degrees of freedom, hence more parameters to be fitted than required, perform on average worse as indicated by the ensembles (Fig. 9B). For the times series with fast gating, we obtained similar results. On the first look, we obtained the same picture as for the time series without fast gating (Fig. 9C). Interestingly, in the zoomed in view (Fig. 9D) it is clear that the OCCO model failed to be distinguished from the input COCO model and the models with more degrees of freedom. Numerical matrix transformation (see Kienker, 1989) revealed that the best solution of the OCCO Markov model is equivalent to the input COCO model whereas we did not find an equivalent solution for the slow gating COCO model (Fig. 9B) within the parameter space. Obviously, the ensemble fits are also suitable to indicate whether an equivalent model exists.

**Figure 9.**
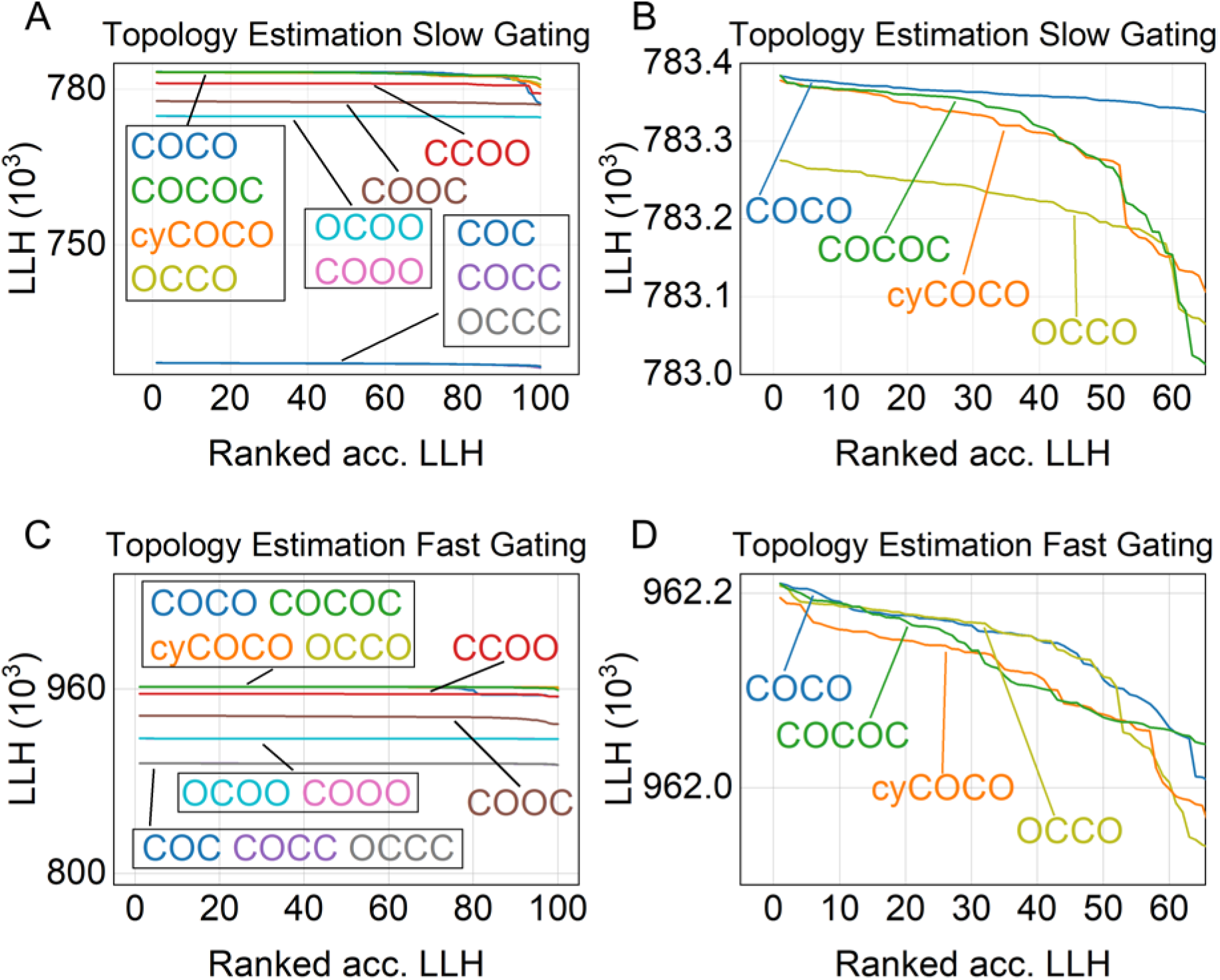
Markov model topology estimation with the 2D-fit. Two Pseudo experimental time series *x_exp_* [*t*] were generated with the following Markov models, one below the frequency of the low-pass filter 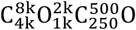 (in Hz, model 1) and another model with two higher rates 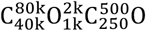 (in Hz, model 2). The length of the time series was 10M samples and SNR = 5. All other settings were according to table 1. *x*_*exp*,1_[*t*] and *x*_*exp*,2_[*t*] were analysed with the 2D-fit with an ensemble size of 100 using different Markov model topologies as depicted in the graphs. The topologies used in this experiment comprised all 4-state models allowing gating between closed and open states, a three state COC model, a cyclic COCO model and the five-state COCOC model. The fit boundaries for all *k_ij_* were 10 Hz to 100 kHz for model (1) and 10 Hz to 1 MHz for model (2). **A** Depicted are the ranked ensemble fits according to the objective score (Eq.2) for all Markov topologies for model 1 and **B** for a zoomed in view of the best performing models. **C** shows the ranked objective score for model 2 and **D** zoomed in to the topologies with the highest score.

## Discussion

Previous implementations of the 2D-Dwell-Time-Fit (2D-fit) suffered from the tremendous computational demanding simulation process of time series (Magleby and Weiss, 1990b), potentiated by the use of a genetic algorithm (Huth et al., 2006). In the previous version of the fit program, single fits took days or even weeks to complete making a thorough modeling of a time series a tedious process. We now solved this issue by implementing a Message Passing Interface (MPI) allowing to distribute time series simulations to worker processes while the manager process governs the genetic algorithm. We have shown, that our implementation allows for massive parallel computing (Fig. 2A). However, using a simple genetic algorithm (Wall, 1996) poses an important limitation. The maximum number of utilizable worker processes is limited by population size of the genetic algorithm. By using multiple MPI processes in parallel the hard limit is set only by the number of available nodes. In addition, we demonstrated that with increasing population size, e.g. for Markov models with higher complexity, the number of processes can be scaled up proportionally without considerably compromising parallel efficiency (Fig. 2B).

With parallel processing, it is now feasible to conduct ensemble fits, which is the most important achievement of this study. In our previous study we have noted that the calculated deviation between the experimental and the simulated 2D-histogram is heavily affected by the stochastic nature of the simulation process (Huth et al., 2006). Conventional fit algorithms therefore fail in the process of optimizing rates of the Markov model as exemplified with the Simplex algorithm. Therefore, we employed a genetic algorithm and demonstrated its superiority (Huth et al., 2006). In line with previous reports we showed (Fig. 3A) that even with the genetic algorithm a large fraction of the ensemble solutions were distant from the global optimum (Magleby and Weiss, 1990b). However, the good news was that by ranking the fits according to the objective score we reached the global maximum in a task of determining rate constants of a five-state COCOC model (Fig. 3A) with several ensemble solutions. In general, this approach allows selecting results that are very likely close to the global maximum and therefore have a high precision of the rates indicated by the MAPE (Fig. 3B).

We then employed the fit ensembles to optimize the settings of the genetic algorithm (table 1) and outlined how to establish a population size of the genetic algorithm and an ensemble size for obtaining reliable results. In addition, we established the limits of the 2D-fit with a particular focus on time series’ noise and gating events beyond the corner frequency of the low-pass filter (fast gating). We showed that at least an SNR of one is required for reliably extracting rate constants of the underlying Markov model. This finding is in accordance with our previous study using the 2D-fit with a simpler CO model (Huth et al., 2006) and is exceptional for patch-clamp analysis. Importantly, we demonstrated that the 2D-fit could still extract fast gating events in case of a low SNR (Fig. 6J). In this setting and with a better SNR of five (Fig. 5D) it is capable to extract rate constants ten times larger than the corner of frequency of the low-pass filter. In our previous study, we obtained a similar result, but we were not able to establish an upper limit (Huth et al., 2006). In addition, we now could establish that in contrast to the fit of β-distributions alone (Fig. 7C,F; Schröder et al., 2005) fast rate estimation is not impaired by additional slow rates present in the model.

Topology estimation of the Hidden Markov model is a key feature of the 2D-fit (Magleby and Weiss, 1990a, 1990b; Huth et al., 2006, 2008). Connecting open and closed intervals in the 2D-histogram preserves the coherence of states and allows deducing the Markov chain (Labarca et al., 1985; Magleby and Weiss, 1990a). This finding is of paramount importance, because the chain can connect the observable time series with the underlying conformational changes of the ion channel protein. In this study we have now extended previous findings by demonstrating that ensemble simulations are mandatory for reliable model discrimination (Fig. 9). Markov Models are not always unique, there exist classes of Markov Models that exhibit the same steady state behavior. In theory they are indistinguishable from one another and are called equivalent models. A transformation to determine equivalent models was proposed in (Kienker, 1989). In practice, thankfully, not all transformed models yield physically meaningful rates (i.e. they are negative or too high). This reduces the number of equivalent models that are relevant. As shown, the ensemble fits allow identifying potential equivalent solutions. In a real experiment, several time series will be analyzed, e.g. obtained at different voltages. By analyzing all time series with probable equivalent Markov chains, in the end it should be possible to identify the matching topology.

All algorithms dealing with modeling of Dwell-Time histograms have to idealize the time series prior hand. Considerable effort was undertaken in recent years to improve event detection beyond simple threshold methods, even without input of baseline and channel conductance. For instance, Celik and colleagues (Celik et al., 2020) used an unsupervised deep neural network to detect single gating events whereas Gnanasambandam and colleagues used an algorithm based on Rissanen’s minimum description length principle (Gnanasambandam et al., 2017). Despite these advances, these studies did not improve the general problems associated with idealization (Blatz and Magleby, 1986; Magleby and Weiss, 1990b; Qin and Li, 2004; reviewd in Qin, 2007). Ultimately, the temporal resolution of the dwell-times is severely limited by the frequency of the low-pass filter and hence fast rates could not be modeled. In addition, in case of fast gating the conducting current level is apparently reduced (Fig. 5B). It is unknown whether the above-mentioned algorithms are capable to deal with this issue. The basic idea of the 2D-fit algorithm is that errors made during the idealization process are canceling out in the experimental and simulated times series (Magleby and Weiss, 1990b; Huth et al., 2006). Based on the results of this study we hypothesize that an accurate idealization is not required at all for proper modeling with the 2D-fit. This assumption is made given the observation that the fitting results are partly independent from the correct setting of the current level, implying improper idealization (Fig. 5E,F; 8F). It would be plausible that an alternative convolution of the time series exists to yield proper models with an even better outcome than the Hinkley idealization detector used here (Hinkley, 1970; Schultze and Draber, 1993; Draber and Schultze, 1994). Regardless of the idealization process, the current level of the conducting state is an important feature of the channel protein. Therefore, automatic level detection as outlined in this study is a powerful feature. Interestingly, using a LLH and current amplitude score together is not compatible (Fig. 7G, 8D) as noted before (Schröder et al., 2005). However, while not generally performing well, the 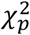 is compatible and improved level detection. In the future, it should allow to investigate if apparent subconductance events in patch-clamp recordings are caused by fast gating or not.

Considerable effort was undertaken to improve the bandwidth of patch-clamp recordings (Levis and Rae, 1993; Parzefall et al., 1998) with some recent advances (Mukhtasimova et al., 2016; Hartel et al., 2018). Equally important, the analysis of fast gating events was improved (Rauh et al., 2017). Here, the capability of the 2D-fit to extract fast gating events on a noisy background is outstanding to maximize recording bandwidth. Given that the 2D-fit can extract rates ten times higher than the cut-off frequency of the low-pass filter, gating events in the range below one μs can be reliably resolved. With this high temporal resolution, a new class of conformational changes in ion channel proteins can be investigated bridging patch-clamp experiments and molecular dynamic simulations.

### Conclusion

In this study, we utilized only synthetic data for fitting purposes. However, it was previously demonstrated that original non-stationary single-channel patch-clamp data could be successfully modeled using the 2D-fit with simulations (Huth et al., 2008). In addition, the fit could be employed in the case of multiple ion channels in one time series (Huth et al., 2006). In summary, employing HPC clusters with ensemble fits overcomes the previous limitations and unhandiness of the 2D-fit by obtaining statistics and employing level detection for a highly automated, standardized and little time consuming approach. Reliable model discrimination and the capability of extracting fast rates on a noisy background makes the 2D-fit stand out among other modeling programs. In addition, beyond the powerful capabilities of the algorithm a huge advantage compared to other algorithms will be the possibility to estimate the modeling quality by statistical means and therefore obtaining a high accuracy.

## Acknowledgements

The authors gratefully acknowledge the scientific support and HPC resources provided by the Erlangen National High Performance Computing Center (NHR@FAU) of the Friedrich-Alexander-Universität Erlangen-Nürnberg (FAU) under the NHR project b109dc. NHR funding is provided by federal and Bavarian state authorities. NHR@FAU hardware is partially funded by the German Research Foundation (DFG) – 440719683. This study was supported by the Deutsche Forschungsgemeinschaft (DFG, German Research Foundation, HU 2358/1-1) to TH. The present work was performed in (partial) fulfillment of the requirements for obtaining the degree Dr. hum. biol.

## References

Albertsen A, Hansen U-P (1994) Estimation of kinetic rate constants from multi-channel recordings by a direct fit of the time series. Biophys J 67:1393–1403.

Benndorf K (1995) Low-Noise Recording. In: Single-Channel Recording, 2nd ed. (Sakmann B, Neher E, eds), pp 129–145. Boston, MA: Springer US.

Blatz a L, Magleby KL (1986) Correcting single channel data for missed events. Biophys J 49:967–980.

Celik N, O’Brien F, Brennan S, Rainbow RD, Dart C, Zheng Y, Coenen F, Barrett-Jolley R (2020) Deep-Channel uses deep neural networks to detect single-molecule events from patch-clamp data. Commun Biol 3:3.

Colquhoun D, Hatton CJ, Hawkes AG (2003) The quality of maximum likelihood estimates of ion channel rate constants. J Physiol 547:699–728.

Colquhoun D, Hawkes AG (1977) Relaxation and Fluctuations of Membrane Currents that Flow through Drug-Operated Channels. J Chem Inf Model 53:1689–1699.

Draber S, Schultze R (1994) Detection of jumps in single-channel data containing subconductance levels. Biophys J 67:1404–1413.

Fredkin DR, Rice JA (1992) Maximum Likelihood Estimation and Identification Directly from Single-Channel Recordings. Proc R Soc B Biol Sci 249:125–132.

Gnanasambandam R, Nielsen MS, Nicolai C, Sachs F, Hofgaard JP, Dreyer JK (2017) Unsupervised Idealization of Ion Channel Recordings by Minimum Description Length: Application to Human PIEZO1-Channels. Front Neuroinform 11:31.

Hamill OP, Marty A, Neher E, Sakmann B, Sigworth FJ (1981) Improved patch-clamp techniques for high-resolution current recording from cells and cell-free membrane patches. Pflügers Arch - Eur J Physiol 391:85–100.

Hartel AJW, Ong P, Schroeder I, Giese MH, Shekar S, Clarke OB, Zalk R, Marks AR, Hendrickson WA, Shepard KL (2018) Single-channel recordings of RyR1 at microsecond resolution in CMOS-suspended membranes. Proc Natl Acad Sci 115:E1789–E1798.

Hinkley D (1970) Inference about the change point from cumulative sum-tests. Biometrika 58:509–523.

Holland JH (1992) Adaptation in Natural and Artificial Systems. Ann Arbor: The MIT Press.

Horn R, Lange K (1983) Estimating kinetic constants from single channel data. Biophys J 43:207–223.

Huth T, Schmidtmayer J, Alzheimer C, Hansen UP (2008) Four-mode gating model of fast inactivation of sodium channel Nav1.2a. Pflügers Arch - Eur J Physiol 457:103–119.

Huth T, Schroeder I, Hansen U-P (2006) The power of two-dimensional dwell-time analysis for model discrimination, temporal resolution, multichannel analysis and level detection. J Membr Biol 214:19–32.

Kienker P (1989) Equivalence of Aggregated Markov Models of Ion-Channel Gating. Proc R Soc B Biol Sci 236:269–309.

Labarca P, Rice JA, Fredkin DR, Montal M (1985) Kinetic analysis of channel gating. Application to the cholinergic receptor channel and the chloride channel from Torpedo californica. Biophys J 47:469–478.

Lampert A, Korngreen A (2014) Markov modeling of ion channels: implications for understanding disease. Prog Mol Biol Transl Sci 123:1–21.

Levis RA, Rae JL (1993) The use of quartz patch pipettes for low noise single channel recording. Biophys J 65:1666–1677.

Magleby KL, Weiss DS (1990a) Identifying Kinetic Gating Mechanisms for Ion Channels by Using Two-Dimensional Distributions of Simulated Dwell Times. Proc R Soc B Biol Sci 241:220–228.

Magleby KL, Weiss DS (1990b) Estimating kinetic parameters for single channels with simulation. A general method that resolves the missed event problem and accounts for noise. Biophys J 58:1411–1426.

Mukhtasimova N, daCosta CJB, Sine SM (2016) Improved resolution of single channel dwell times reveals mechanisms of binding, priming, and gating in muscle AChR. J Gen Physiol 148:43–63.

Neher E, Sakmann B (1976) Single-channel currents recorded from membrane of denervated frog muscle fibres. Nature 260:799–802.

Parzefall F, Wilhelm R, Heckmann M, Dudel J (1998) Single channel currents at six microsecond resolution elicited by acetylcholine in mouse myoballs. J Physiol 512:181–188.

Qin F (2007) Principles of single-channel kinetic analysis. In: Methods in molecular biology (Clifton, N.J.), pp 253–286.

Qin F, Auerbach A, Sachs F (1997) Maximum likelihood estimation of aggregated Markov processes. Proc R Soc B Biol Sci 264:375–383.

Qin F, Auerbach A, Sachs F (2000) Hidden Markov Modeling for Single Channel Kinetics with Filtering and Correlated Noise. Biophys J 79:1928–1944.

Qin F, Li L (2004) Model-based fitting of single-channel dwell-time distributions. Biophys J 87:1657–1671.

Rauh O, Hansen U-P, Mach S, Hartel AJW, Shepard KL, Thiel G, Schroeder I (2017) Extended beta distributions open the access to fast gating in bilayer experiments-assigning the voltage-dependent gating to the selectivity filter. FEBS Lett 591:3850–3860.

Rechenberg I (1978) Evolutionsstrategien. In, pp 83–114. Stuttgart: Fromman-Hozboog Verlag.

Schröder I, Harlfinger P, Huth T, Hansen UP (2005) A subsequent fit of time series and amplitude histogram of patch-clamp records reveals rate constants up to 1 per microsecond. J Membr Biol 203:83–99.

Schroeder I (2015) How to resolve microsecond current fluctuations in single ion channels: The power of beta distributions. Channels 9:262–280.

Schultze R, Draber S (1993) A nonlinear filter algorithm for the detection of jumps in patch-clamp data. J Membr Biol 132:41–52.

Venkataramanan L, Sigworth FJ (2002) Applying hidden Markov models to the analysis of single ion channel activity. Biophys J 82:1930–1942.

Wall MB (1996) A genetic algorithm for resource-constrained scheduling.

